# Cell cycle perturbation uncouples mitotic progression and invasive behavior in a post-mitotic cell

**DOI:** 10.1101/2023.03.16.533034

**Authors:** Michael A. Q. Martinez, Chris Z. Zhao, Frances E. Q. Moore, Callista Yee, Wan Zhang, Kang Shen, Benjamin L. Martin, David Q. Matus

## Abstract

The acquisition of the post-mitotic state is crucial for the execution of many terminally differentiated cell behaviors during organismal development. However, the mechanisms that maintain the post-mitotic state in this context remain poorly understood. To gain insight into these mechanisms, we used the genetically and visually accessible model of *C. elegans* anchor cell (AC) invasion into the vulval epithelium. The AC is a terminally differentiated uterine cell that normally exits the cell cycle and enters a post-mitotic state, initiating contact between the uterus and vulva through a cell invasion event. Here, we set out to identify the set of negative cell cycle regulators that maintain the AC in this post-mitotic, invasive state. Our findings revealed a critical role for CKI-1 (p21^CIP1^/p27^KIP1^) in redundantly maintaining the post-mitotic state of the AC, as loss of CKI-1 in combination with other negative cell cycle regulators—including CKI-2 (p21^CIP1^/p27^KIP1^), LIN-35 (pRb/p107/p130), FZR-1 (Cdh1/Hct1), and LIN-23 (β-TrCP)—resulted in proliferating ACs. Remarkably, time-lapse imaging revealed that these ACs retain their ability to invade. Upon examination of a node in the gene regulatory network controlling AC invasion, we determined that proliferating, invasive ACs do so by maintaining aspects of pro-invasive gene expression. We therefore report that the requirement for a post-mitotic state for invasive cell behavior can be bypassed following direct cell cycle perturbation.

## INTRODUCTION

The coordination of terminal differentiation and cell cycle exit is critical for tissue morphogenesis during organismal development. The general model for how terminally differentiated cells exit the cell cycle involves the repression of cyclin/cyclin-dependent kinase (CDK) activity by negative cell cycle regulators. These regulators include the CDK inhibitors (CKIs), retinoblastoma (Rb) family members, and E3 ubiquitin ligase complexes, such as the anaphase-promoting complex/cyclosome (APC/C) and the Skp-Cullin-F-box protein (SCF) complex (Buttitta and Edgar, 2007; Ruijtenberg and van den Heuvel, 2016). However, despite the generally accepted model, it remains unclear how terminally differentiated cells maintain their post-mitotic state once they have exited the cell cycle.

Cell cycle exit following terminal differentiation is considered critical for tissue morphogenesis because it allows for the execution of specialized cell behaviors. According to the “go versus grow” dichotomy, a mutually exclusive relationship exists between these cell behaviors and cell proliferation (Kohrman and Matus, 2017). However, cell cycle exit and cell morphogenetic behaviors are not always functionally linked. For example, neural crest delamination, a specialized epithelial-to-mesenchymal transition (EMT), occurs in a subset of neural crest cells at the G1/S boundary (Burstyn-Cohen and Kalcheim, 2002). Additionally, it is possible to experimentally uncouple morphogenetic cell behavior from cell cycle exit. In the developing mouse retina, genetic ablation of Rb paralogs leads to fully differentiated neurons that retain their ability to form neurites and synapses while proliferating (Ajioka et al., 2007). Similar results have been observed in mouse inner ear hair cells following the loss of Rb (Sage et al., 2005).

Furthermore, recent evidence from two-dimensional and three-dimensional *in vitro* assays using skin cancer cells has found that these cells can simultaneously invade and proliferate (Haass et al., 2014; Vittadello et al., 2020). Together, these findings suggest that the terminal differentiation programs that drive key morphogenetic cell behaviors can occur without exiting the cell cycle.

Prior research from our laboratory and others has utilized a simple *in vivo* model of developmental cell invasion, *C. elegans* anchor cell (AC) invasion, to functionally dissect the relationship between cell cycle exit and morphogenetic cell behavior (Deng et al., 2020; Matus et al., 2015; Medwig-Kinney et al., 2020; Smith et al., 2022). During the third larval stage of post-embryonic nematode development, the terminally differentiated and post-mitotic AC initiates the uterine-vulval connection by breaching the basement membrane (BM) that separates these two tissues (Sherwood and Sternberg, 2003) (Fig. 1A, left). AC invasion occurs in synchrony with the divisions of P6.p, the underlying 1°-fated vulval precursor cell (VPC), such that by the P6.p 4-cell stage, BM removal is complete (Sherwood and Sternberg, 2003). Robust control of the AC cell cycle appears to be required for invasive differentiation, as loss of the post-mitotic state leads to a complete loss of invasive capacity (Matus et al., 2015) (Fig. 1A, right). The SWI/SNF chromatin remodeling complex, specifically the evolutionarily conserved BAF assembly, along with three conserved transcription factors—EGL-43 (EVI1), HLH-2 (E/Daughterless), and NHR-67 (TLX/Tailless)—function as key regulators of AC invasion by maintaining the AC in a post-mitotic state (Deng et al., 2020; Matus et al., 2015; Medwig-Kinney et al., 2020; Smith et al., 2022). Experimental evidence supports the relationship between cell cycle exit and AC invasion, as AC-specific expression of the CDK inhibitor CKI-1 (p21^CIP1^/p27^KIP1^) following the loss of NHR-67 completely rescues BM invasion (Matus et al., 2015). Conversely, CKI-1 knockdown only rarely induces AC proliferation (Matus et al., 2015). The requirement for the loss of multiple negative regulators for cell cycle re-entry has been demonstrated in other terminally differentiated cell types (Buttitta and Edgar, 2007). Therefore, it is possible that additional negative regulators of the AC cell cycle redundantly contribute to maintaining the AC in a post-mitotic, invasive state.

**Figure 1.**
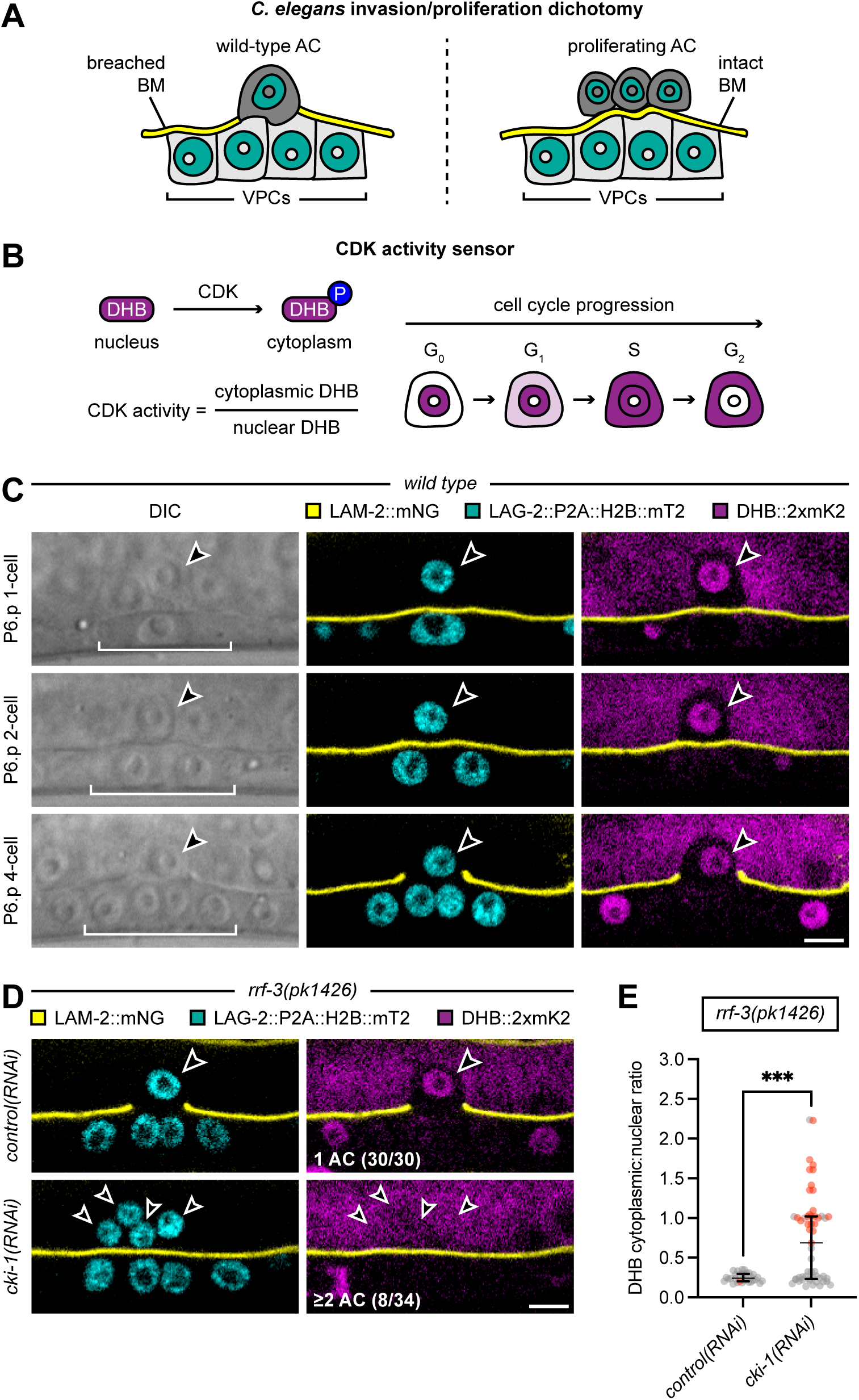
A moderate increase in AC proliferation is observed upon systemic depletion of CKI-1. (A) Schematic of the *C. elegans* invasion/proliferation dichotomy. (B) Schematic of the CDK activity sensor, DHB, which is phosphorylated by CDK and translocates from the nucleus to the cytoplasm in response to CDK activity during cell cycle progression. The equation used to quantify CDK activity is shown. (C) DIC (left) and confocal images merging LAM-2::mNeonGreen with LAG-2::P2A::H2B::mTurquoise2 (middle) and DHB::2xmKate2 (right) from the P6.p 1-cell stage to the P6.p 4-cell stage are shown. Arrowheads and brackets indicate the position of the AC and P6.p cells, respectively. The scale bar represents 5 µm. (D) Confocal images merging LAM-2::mNeonGreen with LAG-2::P2A::H2B::mTurquoise2 (left) and DHB::2xmKate2 (right) from the P6.p 4-cell stage are shown. Arrowheads indicate the position of the AC(s) in *rrf-3(pk1426)* animals following *control(RNAi)* and *cki-1(RNAi)* treatment. The scale bar represents 5 µm. (E) Scatter plot displays the median and IQR of DHB::2xmKate2 ratios in the AC(s) of *rrf-3(pk1426)* animals following *control(RNAi)* and *cki-1(RNAi)* treatment (N ≥ 30 animals per treatment). Gray dots correspond to cells where the BM is broken, and red dots correspond to cells where the BM is intact. Statistical significance was determined by a Mann-Whitney test (P < 0.0004). See also Figure S1, S2.

Here, we used *C. elegans* AC invasion as a genetically dissectible *in vivo* model of developmental cell invasion to investigate the relationship between cell cycle regulation and cell invasive behavior. Previous data supported a model where the AC must exit the cell cycle and enter a post-mitotic state to acquire an invasive phenotype (Deng et al., 2020; Matus et al., 2015; Medwig-Kinney et al., 2020; Smith et al., 2022). Our objective was to identify the specific set of negative cell cycle regulators that maintain the AC in a post-mitotic state. By combining a live CDK activity sensor with enhanced RNAi and available null or hypomorphic alleles, we identified a crucial role for CKI-1 in redundantly maintaining the post-mitotic state of the AC. Through time-lapse imaging, we observed that the loss of CKI-1 in combination with another negative cell cycle regulator resulted in proliferating ACs that retained their invasive abilities. Using two reporters of the gene regulatory network that controls AC invasion, we demonstrated that the loss of multiple negative cell cycle regulators led to proliferating, invasive ACs that maintained some aspects of pro-invasive gene expression. Thus, our work demonstrates that in the AC, invasive cell behavior can be uncoupled from cell cycle exit.

## RESULTS

### Improved efficiency of CKI-1 depletion increases AC proliferation

In our previous attempt to deplete CKI-1 through RNAi, we employed a tissue-specific approach by reintroducing RDE-1 into the uterine cells of *rrf-3(pk1426); rde-1(ne219)* double mutants using the *fos-1a* promoter (Hagedorn et al., 2009; Matus et al., 2010; Morrissey et al., 2014; Qadota et al., 2007). In this uterine-specific RNAi-sensitive background, the loss of either the pro-invasive transcription factor EGL-43 or NHR-67 led to AC proliferation without BM invasion in over 60% of animals (Matus et al., 2015; Medwig-Kinney et al., 2020). Conversely, uterine-specific RNAi-mediated depletion of CKI-1, a canonical negative regulator of the *C. elegans* cell cycle (Feng et al., 1999; Fukuyama et al., 2003; Hong et al., 1998), rarely resulted in this phenotype, occurring in about 3% of animals (Matus et al., 2015). For interpreting both these and subsequent results, it is important to note that the wild-type AC is born into a CDK-low, post-mitotic state, invades 100% of the time, and never re-enters the cell cycle (Matus et al., 2015; Sherwood and Sternberg, 2003; Smith et al., 2022). Given the low occurrence of proliferating ACs following CKI-1 depletion, we hypothesized that this lack of phenotype could be attributed to the functional redundancy of CKI-1 with other negative cell cycle regulators and/or inefficient CKI-1 depletion.

First, we aimed to enhance the efficiency of *cki-1(RNAi)*. Building upon our previous successes with the highly efficient RNAi vector, T444T (Martinez et al., 2020; Medwig-Kinney et al., 2020; Smith et al., 2022), we generated a new RNAi construct for *cki-1*. Unlike the standard RNAi vector, L4440 (Timmons and Fire, 1998), T444T includes a pair of T7 terminator sequences to prevent transcription of the vector backbone (Sturm et al., 2018). Subsequently, we evaluated the efficacy of the T444T-based *cki-1(RNAi)* construct in the uterine-specific RNAi strain (*rrf-3(pk1426); rde-1(ne219); fos-1ap::rde-1*), utilizing fluorescent reporters to visualize the AC cell membrane and BM. Among the animals exposed to this construct, 5% had ≥2 ACs (Fig. S1A), representing a 2% increase in the proliferative AC phenotype compared to the original *cki-1(RNAi)* experiment (Matus et al., 2015). Notably, whenever multiple ACs were observed, there was consistently an accompanying BM gap. We hypothesized that this finding could be attributed to either a delayed RNAi effect or the reestablishment of the post-mitotic, invasive state following AC proliferation. Alternatively, this result raises the intriguing possibility that ACs can simultaneously invade and proliferate.

### Systemic depletion of CKI-1 causes an increase in AC proliferation

To better visualize the interplay between AC invasion and proliferation, we generated a three-color imaging strain for simultaneous monitoring of the AC, cell cycle, and BM (Fig. 1A-C). First, to visualize the AC nucleus (shown in cyan), we used an endogenous transcriptional reporter of *lag-*2 (*Delta*), LAG-2::P2A::H2B::mTurquoise2 (Medwig-Kinney et al., 2022). To visualize and quantify cell cycle state (shown in red), we used a ubiquitous CDK activity sensor, DHB::2xmKate2 (Adikes et al., 2020). Finally, to accurately assess BM invasion (shown in yellow), we used an endogenous translational reporter of *lam-2*, LAM-2::mNeonGreen, to label laminin, a core BM component (Jayadev et al., 2019). Conveniently, the histone-based *lag-2* transcriptional reporter also functions as a P6.p lineage tracker, facilitating the developmental staging of AC invasion (Fig. 1A,C). The CDK activity sensor is a fluorescent fragment of human DNA helicase B (DHB), which is predominantly nuclear in post-mitotic cells, such as the wild-type AC, and becomes increasingly cytoplasmic in proliferating cells as they progress through the cell cycle (Fig. 1B,C). Thus, the cytoplasmic-to-nuclear ratio of DHB fluorescence provides a visual and quantitative readout of CDK activity, and more generally, cell cycle progression (Adikes et al., 2020; Martinez and Matus, 2022; Spencer et al., 2013; van Rijnberk et al., 2017).

With the three-color imaging strain at our disposal, our initial goal was to assess whether our enhanced *cki-1(RNAi)* construct could successfully induce AC cell cycle entry in a wild-type background. As expected, 100% of *control(RNAi)* animals showed a post-mitotic, invasive AC (Fig. S1B,C; Table 1). Unexpectedly, we observed the same results for *cki-1(RNAi)*, with all animals possessing a single post-mitotic, invasive AC (Fig. S1B,C; Table 1). We next introduced the *rrf-3(pk1426)* null allele (Simmer et al., 2002), which on its own confers RNAi hypersensitivity in many somatic tissues, including the uterus (Matus et al., 2010). While all *rrf-3(pk1426); control(RNAi)* animals contained a post-mitotic AC, 24% of *rrf-3(pk1426); cki-1(RNAi)* animals possessed ≥2 ACs (Fig. 1D,E; Table 1). In this RNAi-hypersensitive background, CKI-1 depletion correlated with a significant increase in AC-specific CDK activity when compared to control animals (0.25 ± 0.06 (n = 30) vs. 0.73 ± 0.56 (n = 55)). Notably, in 3 of the 8 animals with ≥2 ACs, BM invasion was observed (Table 1). Taken together, these data demonstrate that systemic depletion of CKI-1, via the use of an enhanced RNAi construct in an RNAi-hypersensitive background, can induce the AC to re-enter the cell cycle.

**Table 1.** Genetic analysis of negative cell cycle regulators during AC invasion.

|  |  | % invasion |  | % no invasion |  |  |
| --- | --- | --- | --- | --- | --- | --- |
| **Genotype** | **Treatment** | **1 AC** | **≥2 AC** | **1 AC** | **≥2 AC** | **N** |
| *wild type* | *control(RNAi)* | 100 | 0 | 0 | 0 | 27 |
| *rrf-3(pk1426)* |  | 97 | 0 | 3 | 0 | 30 |
| *cdc-14(he141)* |  | 100 | 0 | 0 | 0 | 35 |
| *cki-2(ok2105)* |  | 100 | 0 | 0 | 0 | 30 |
| *lin-35(n745)* |  | 100 | 0 | 0 | 0 | 31 |
| *fzr-1(ku298)* |  | 100 | 0 | 0 | 0 | 32 |
| *lin-23(e1883)* |  | 100 | 0 | 0 | 0 | 34 |
| *wild type* | *cki-1(RNAi)* | 100 | 0 | 0 | 0 | 29 |
| *rrf-3(pk1426)* |  | 76 | 9 | 0 | 15 | 34 |
| *cdc-14(he141)* |  | 100 | 0 | 0 | 0 | 35 |
| *cki-2(ok2105)* |  | 87 | 13 | 0 | 0 | 31 |
| *lin-35(n745)* |  | 85 | 12 | 0 | 3 | 33 |
| *fzr-1(ku298)* |  | 65 | 29 | 0 | 6 | 34 |
| *lin-23(e1883)* |  | 19 | 74 | 0 | 7 | 31 |

AC invasion was scored at the P6.p 4-cell stage of vulval development, the stage when AC invasion normally occurs, using AC/VPC, BM, and cell cycle fluorescent reporters.

Unfortunately, we were unable to assess the effect of the *cki-1(gk132)* null allele on AC proliferation (The *C. elegans* Deletion Mutant Consortium, 2012), as it leads to embryonic lethality in 100% of homozygotes (Buck et al., 2009) (Fig. S2A). Instead, we crossed the *cdc-14(he141)* null allele into the three-color imaging strain (Saito et al., 2004). *cdc-14* encodes a dual-specificity phosphatase that positively regulates CKI-1 activity in a number of larval tissues (Clayton et al., 2008; Roy et al., 2011; Roy et al., 2014; Saito et al., 2004), likely by preventing its phosphorylation-dependent degradation (Saito et al., 2004). Nonetheless, 100% of *cdc-14(he141)* animals had a post-mitotic, invasive AC, even in the presence of *cki-1(RNAi)* (Fig. S2B,C; Table 1), suggesting that CDC-14 is either not a regulator of CKI-1 in the AC or that its function is redundant with that of another phosphatase.

### CKI-1 is critical for maintaining the post-mitotic state of the AC

In other model systems, the post-mitotic state of terminally differentiated cells is maintained through functional redundancy among negative cell cycle regulators (Buttitta et al., 2007; Ma et al., 2019). Thus far, our data demonstrate that depletion of CKI-1 alone can induce the AC to re-enter the cell cycle, albeit with incomplete penetrance. We therefore hypothesized that this incomplete penetrance may be attributed to either partial depletion of CKI-1 or compensatory mechanisms involving other negative cell cycle regulators, as recently demonstrated in the developing *C. elegans* vulva (Portegijs, 2019).

Only a handful of factors negatively regulate the somatic cell cycle during *C. elegans* larval development (Kipreos and van den Heuvel, 2019). These include the CDK inhibitors CKI-1 and CKI-2 (p21^CIP1^/p27^KIP1^) (Buck et al., 2009; Hong et al., 1998), the sole Rb family member LIN-35 (pRb/p107/p130) (Boxem and van den Heuvel, 2001; Boxem and van den Heuvel, 2002), the APC/C co-activator FZR-1 (Cdh1/Hct1) (Fay et al., 2002), and the F-box/WD-repeat protein LIN-23 (β-TrCP) (Kipreos et al., 1996; Kipreos et al., 2000; Nayak et al., 2002). Although they all function during normal cell cycle exit, their role in maintaining the post-mitotic state is poorly understood.

To determine their role in the AC, we generated four separate three-color imaging strains. Three of these strains include putative null alleles, i.e., *cki-2(ok2105)*, *lin-35(n745)*, or *lin-23(e1883)* (Kipreos et al., 2000; Lu and Horvitz, 1998; The *C. elegans* Deletion Mutant Consortium, 2012). The fourth strain harbors *fzr-1(ku298)*, a hypomorphic allele, as FZR-1 function is required for viability (Fay et al., 2002). However, regardless of their genotype, live imaging and subsequent quantification of CDK activity demonstrated that they all possessed a post-mitotic, invasive AC (Fig. 2; Table 1). These results were consistent with what we found after depleting CKI-1 by RNAi in a wild-type background of the three-color imaging strain (Fig. S1B,C; Table 1).

**Figure 2.**
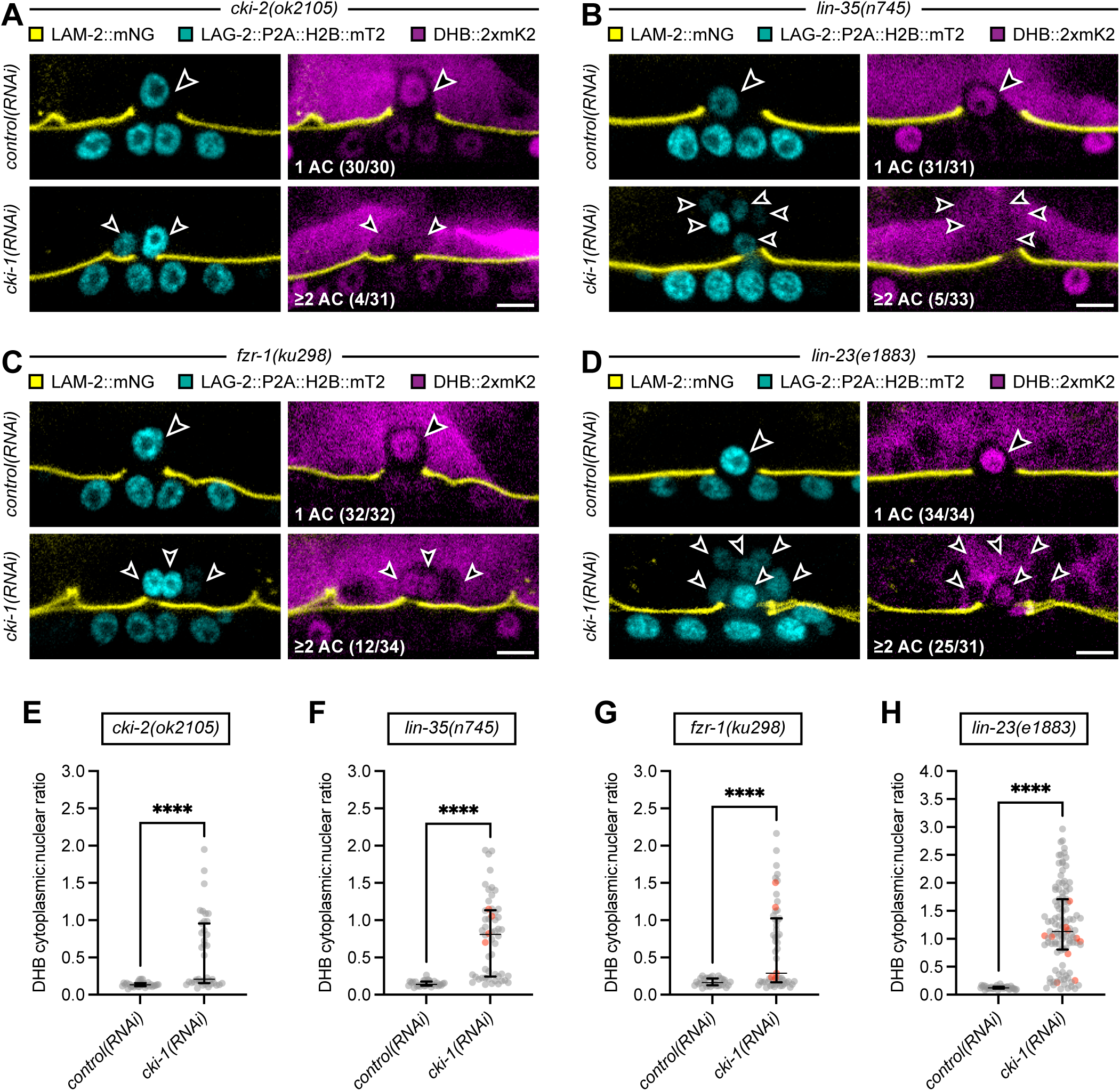
CKI-1 is critical for maintaining the post-mitotic state of the AC. (A-D) Confocal images merging LAM-2::mNeonGreen with LAG-2::P2A::H2B::mTurquoise2 (left) and DHB::2xmKate2 (right) from the P6.p 4-cell stage are shown. Arrowheads indicate the position of the AC(s) in *cki-2(ok2105)* (A), *lin-35(n745)* (B), *fzr-1(ku298)* (C), and *lin-23(e1883)* (D) animals following *control(RNAi)* and *cki-1(RNAi)* treatment. Scale bars represent 5 µm. (E-H) Scatter plots display the median and IQR of DHB::2xmKate2 ratios in the AC(s) of *cki-2(ok2105)* (E), *lin-35(n745)* (F), *fzr-1(ku298)* (G), and *lin-23(e1883)* (H) animals following *control(RNAi)* and *cki-1(RNAi)* treatment (N ≥ 30 animals per treatment). Gray dots correspond to cells where the BM is broken, and red dots correspond to cells where the BM is intact. Statistical significance was determined by a Mann-Whitney test (P < 0.0001).

We next explored whether we could induce AC proliferation by co-depleting CKI-1 in each mutant background. Following *cki-1(RNAi)* treatment, we observed ≥2 ACs in 13%, 15%, 35%, and 81% of *cki-2(ok2105)*, *lin-35(n745)*, *fzr-1(ku298)*, and *lin-23(e1883)* animals, respectively (Fig. 2A-D). Compared to mutant controls, this corresponded to a significant increase in AC-specific CDK activity (0.14 ± 0.03 (n = 30) vs. 0.55 ± 0.50 (n = 39), 0.15 ± 0.04 (n = 31) vs. 0.77 ± 0.52 (n = 54), 0.17 ± 0.05 (n = 32) vs. 0.64 ± 0.57 (n = 55), and 0.13 ± 0.03 (n = 34) vs. 1.22 ± 0.72 (n = 111), respectively (Fig. 2E-H), and this increase was observed in animals with either single or multiple ACs. Thus, direct perturbation of the cell cycle can induce AC cell cycle entry, either through the loss of CKI-1 in an RNAi-hypersensitive background (Fig. 1D,E; Fig. S1A), in combination with null alleles of *cki-2*, *lin-35*, or *lin-23*, or in combination with a hypomorphic allele of *fzr-1*. Together, these findings reveal a critical role for CKI-1 in redundantly maintaining the post-mitotic state of the AC.

### Maintenance of the post-mitotic state is not required for AC invasion

The loss of multiple negative cell cycle regulators not only induced AC proliferation but also resulted in mitotic ACs with an associated underlying gap in the BM (Table 1). This observation suggests that these cycling ACs retained the capacity to invade, which is significant for several reasons. First, the wild-type AC never re-enters the cell cycle (Matus et al., 2015; Smith et al., 2022). Second, the absence of key regulators of AC invasion leads to proliferating ACs that are no longer invasive (Deng et al., 2020; Medwig-Kinney et al., 2020; Smith et al., 2022). Therefore, the identification of cycling, invasive ACs following direct cell cycle perturbation represents a distinct third phenotypic state. However, it is possible that these ACs breach the BM prior to cell cycle entry or after completing mitosis when they return to a post-mitotic state, maintaining the relationship between cell cycle exit and cell invasion.

To further investigate the lack of AC invasion defects following robust cell cycle induction (Table 1), we collected time-lapse movies of proliferating ACs from pre-invasion to post-invasion using a variant of the three-color imaging strain (Fig. 3; Fig. S3). This strain includes an AC-specific F-actin probe, *cdh-3p*::mCherry::moesinABD, allowing us to visualize dynamics of the F-actin cytoskeleton before and during invasion. Importantly, an invasive AC generates protrusions rich in F-actin prior to breaching the BM (Hagedorn et al., 2013; Naegeli et al., 2017). We co-expressed this AC reporter with a ubiquitous CDK activity sensor, DHB::2xmTurquoise2, and an endogenous *lam-2* translational reporter, LAM-2::mNeonGreen. For these experiments, we chose to combine *cki-1(RNAi)* treatment with the *fzr-1(ku298)* hypomorphic allele rather than the *lin-23(e1883)* null allele. In comparison to *lin-23(e1883)* null mutants, which had the highest penetrance of the proliferative AC phenotype in the presence of *cki-1(RNAi)* (Fig. 2D,H; Table 1), the *fzr-1(ku298)* hypomorphs were easier to propagate for the purpose of time-lapse imaging and had the second-highest penetrance of the proliferative AC phenotype under the same treatment conditions (Fig. 2C,G; Table 1).

**Figure 3.**
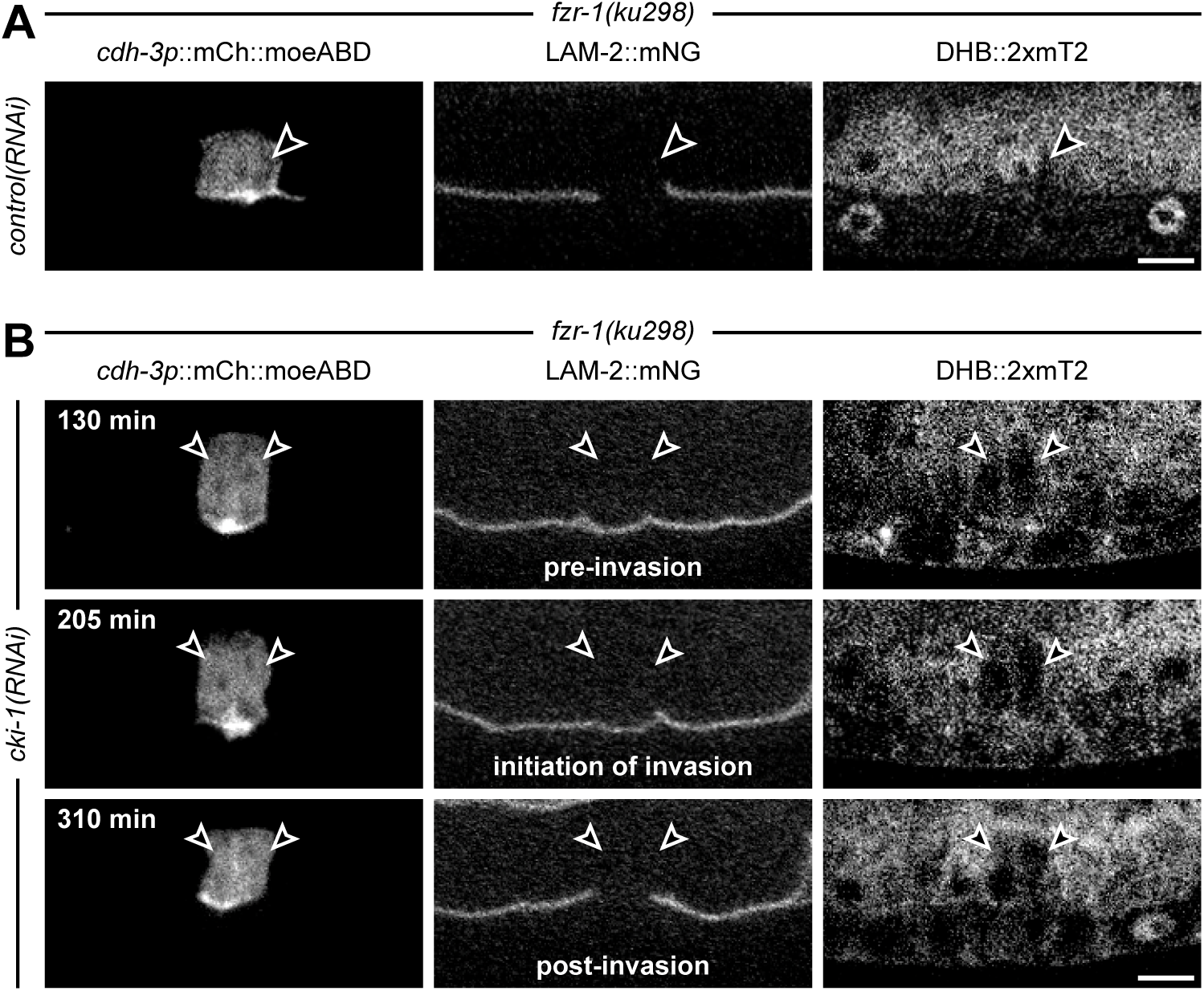
Maintenance of the post-mitotic state is not required for AC invasion. (A) Static images of *cdh-3p*::mCherry::moesinABD (left), LAM-2::mNeonGreen (middle), and DHB::2xmTurquoise2 (right) from the P6.p 4-cell stage are shown. Arrowheads indicate the position of the AC in a *fzr-1(ku298); control(RNAi)* animal. The DHB::2xmTurquoise2 ratio of the AC is 0.26. The scale bar represents 5 µm. (B) Time-lapse images of *cdh-3p*::mCherry::moesinABD (left), LAM-2::mNeonGreen (middle), and DHB::2xmTurquoise2 (right) from pre-AC invasion to post-AC invasion are shown. Arrowheads indicate the position of multiple invading ACs in a *fzr-1(ku298); cki-1(RNAi)* animal. At the initiation of invasion time-point, the DHB::2xmTurquoise2 ratio is 1.56 for the AC on the left and 1.76 for the AC on the right, values that are most consistent with the G2 phase of the cell cycle (Adikes et al., 2020). The scale bar represents 5 µm. See also Movie 1 and Figure S3.

Prior to time-lapse imaging, we captured static images of the invasive AC in *fzr-1(ku298); control(RNAi)* animals (Fig. 3A), which allowed us to obtain a baseline level of CDK activity values (0.31 ± 0.05 (n = 23)). This baseline allowed us to selectively record the activity of pre-invasive ACs in *fzr-1(ku298); cki-1(RNAi)* animals that had already started cycling. Using this method as a screen for proliferating ACs, we were able to collect five time-lapse datasets in which single or multiple cycling, pre-invasive ACs showed an enrichment of the F-actin probe at the invasive membrane (Hagedorn et al., 2013) (1.74 ± 0.36 (n = 7)), followed by successful invasion of the underlying BM (Fig. 3B; Movie 1). In one of these datasets, we observed the division of a pre-invasive AC, with both daughter cells re-entering the cell cycle before invading (Fig. S3). Altogether, these observations led us to conclude that direct cell cycle perturbation results in ACs that can proliferate and invade.

### Aspects of pro-invasive differentiation are partially retained in proliferating, invasive ACs

After finding that direct disruption of the cell cycle led to proliferating, invasive ACs, we investigated whether these ACs maintain a functional pro-invasive gene regulatory network (GRN) (Deng et al., 2020; Medwig-Kinney et al., 2020). This GRN consists of a cell cycle-dependent and cell cycle-independent arm. We hypothesized that gene expression in both arms would remain unchanged in proliferating, invasive ACs following direct cell cycle perturbation.

To test our hypothesis, we quantified endogenous levels of NHR-67, the distal transcription factor in the cell cycle-dependent arm that is upstream of CKI-1. Additionally, we quantified transcriptional levels of the matrix metalloproteinase (MMP) gene, *zmp-1*, a downstream effector of the pro-invasive GRN. Although MMPs are not essential for AC invasion (Kelley et al., 2019), their expression serves as a sensitive readout of invasive differentiation, as disruption of either the cell cycle-dependent or cell cycle-independent arms leads to a near complete loss of MMP expression (Hwang et al., 2007; Matus et al., 2015; Medwig-Kinney et al., 2020; Rimann and Hajnal, 2007; Sherwood et al., 2005). We quantified these levels in two separate *fzr-1(ku298)* strains, each carrying most, if not all, of the original three-color imaging alleles in the presence of *cki-1(RNAi)*. In one strain, we inserted an mNeonGreen tag into the C-terminus of the endogenous *nhr-67* locus using CRISPR/Cas9 (Paix et al., 2015). In the other strain, we replaced the endogenous *lag-2* transcriptional reporter, LAG-2::P2A::H2B::mTurquoise2, with a *zmp-1* transcriptional reporter transgene, *zmp-1p*::CFP (Inoue et al., 2002). Supporting our hypothesis, NHR-67::mNeonGreen expression decreased by only 17% in proliferating, invasive ACs (Fig. 4A,B). Interestingly, *zmp-1p*::CFP expression decreased more significantly by 73% in proliferating, invasive ACs (Fig. 4C,D). These data suggest that while the upstream transcriptional networks that promote AC invasion are intact in mitotic, invasive ACs, downstream effectors exhibit alterations in their expression. Altogether, these results demonstrate that direct cell cycle perturbation uncouples the relationship between cell cycle exit and invasive cellular differentiation.

**Figure 4.**
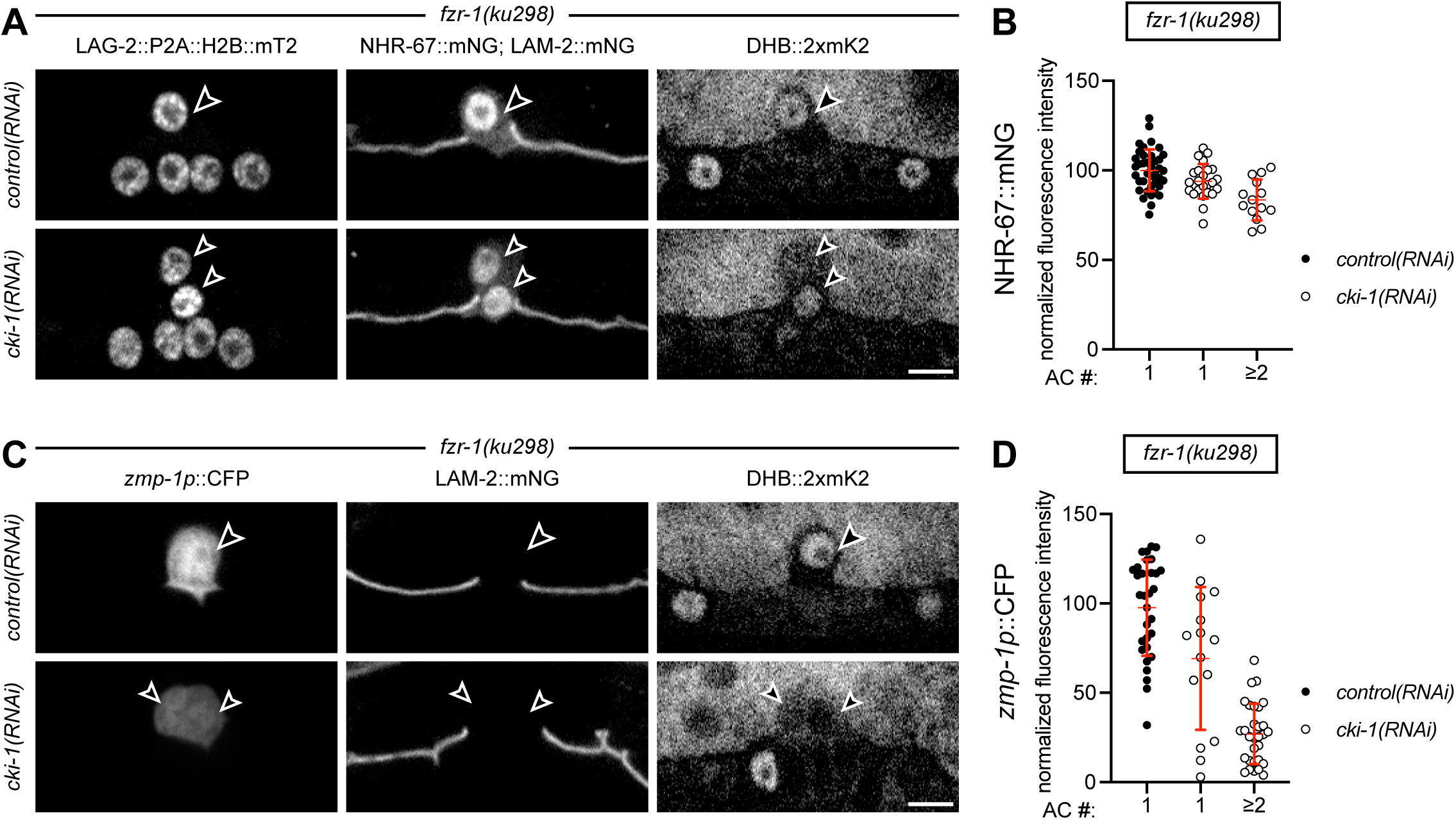
Proliferating, invasive ACs retain a functionally intact pro-invasive GRN. (A) Confocal images of LAG-2::P2A::H2B::mTurquoise2 (left), NHR-67::mNeonGreen (middle), LAM-2::mNeonGreen (middle), and DHB::2xmKate2 (right) from the P6.p 4-cell stage are shown. Arrowheads indicate the position of the AC(s) in *fzr-1(ku298)* animals following *control(RNAi)* and *cki-1(RNAi)* treatment. The scale bar represents 5 µm. (B) Plot displays mean and SD of normalized NHR-67::mNeonGreen intensity in the AC(s) of *fzr-1(ku298)* animals following *control(RNAi)* and *cki-1(RNAi)* treatment (N ≥ 30 animals per treatment). (C) Confocal images of *zmp-1p*::CFP (left), LAM-2::mNeonGreen (middle), and DHB::2xmKate2 (right) from the P6.p 4-cell stage are shown. Arrowheads indicate the position of the AC(s) in *fzr-1(ku298)* animals following *control(RNAi)* and *cki-1(RNAi)* treatment. The scale bar represents 5 µm. (D) Plot displays mean and SD of normalized *zmp-1p*::CFP intensity in the AC(s) of *fzr-1(ku298)* animals following *control(RNAi)* and *cki-1(RNAi)* treatment (N ≥ 31 animals per treatment).

## Discussion

In this study, we characterized the set of negative cell cycle regulators that maintain the post-mitotic state of the invasive *C. elegans* AC. Many terminally differentiated cell types employ a redundant set of negative cell cycle regulators to prevent cell cycle entry (Buttitta et al., 2007; Ma et al., 2019). Similarly, we demonstrate that CKI-1 depletion, in combination with strong loss-of-function alleles of *cki-2*, *lin-35*, *fzr-1*, or *lin-23*, leads to inappropriate AC cell cycle entry. Furthermore, unlike previous perturbations that affected the AC GRN, resulting in proliferating ACs that were unable to invade (Deng et al., 2020; Matus et al., 2015; Medwig-Kinney et al., 2020; Smith et al., 2022), our findings demonstrate that direct targeting of the AC cell cycle machinery generates a population of ACs capable of both proliferating and invading.

We also establish a critical role for CKI-1 in maintaining the post-mitotic state of the AC, as depletion of CKI-1 using an enhanced RNAi construct in a RNAi-hypersensitive background can trigger AC cell cycle entry. In contrast, we show that the AC never re-enters the cell cycle in strong loss-of-function alleles of the other negative regulators of the cell cycle. In the developing *C. elegans* vulva, strong compensatory mechanisms increase the levels of CKIs following the loss of single negative cell cycle regulators (Portegijs, 2019; The et al., 2015). This same compensatory relationship may exist in the AC, explaining the enhanced penetrance of the proliferative AC phenotype following RNAi-mediated depletion of CKI-1 in *cki-2*, *lin-35*, *fzr-1*, and *lin-23* mutant backgrounds (Fig. 2). Unfortunately, GFP tags of the endogenous *cki-1* locus result in a hypermorphic AC phenotype, likely due to an overstabilization of CKI-1 protein (Adikes et al., 2020). Thus, examining endogenous CKI-1 levels in these mutant backgrounds will be challenging. Nevertheless, despite this technical hurdle, the most parsimonious explanation of our results is a model in which the sole loss of other negative cell cycle regulators leads to an upregulation of CKI-1 to maintain a post-mitotic AC (Fig. 5). Therefore, depletion of CKI-1 in these mutant backgrounds relieves this compensatory mechanism, resulting in proliferating ACs that, at least in a subset of cycling ACs, retain the ability to invade (Fig. 5).

**Figure 5.**
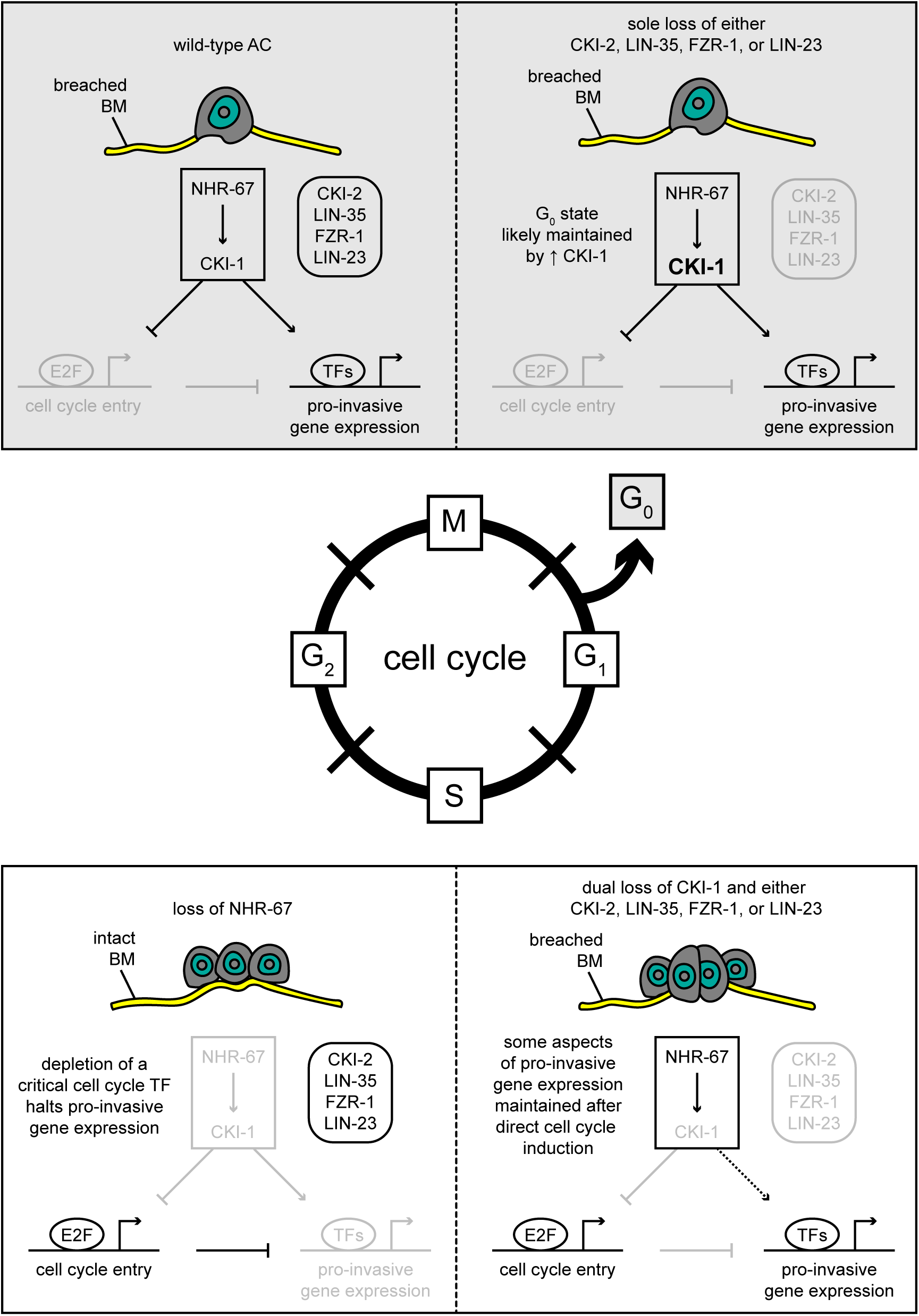
A model for breaking the invasive-proliferative dichotomy. Schematic depicts a summary of previously reported data (left top and bottom) and data from this study (right top and bottom). Wild-type single ACs can only invade in a G0 state (top left, grey box). Loss of single negative cell cycle regulators fail to trigger cell cycle entry, the AC remains in a G0 state, and still invades (top right, grey box). We hypothesize that CKI-1 levels increase to prevent cell cycle entry in response to loss of the other negative cell cycle regulators. Loss of the transcription factor, NHR-67, results in cycling ACs that proliferate and do not invade (bottom left, white box). Dual loss of cell cycle regulators results in proliferative ACs that retain some invasive capacity (bottom right, white box), likely due to core components of the AC invasion GRN remaining functional.

In other developmental contexts, similar observations have been made where cells proceed with their terminal differentiation programs despite re-entering the cell cycle. For example, during *Drosophila* development, the ectopic co-expression of E2F and G1 cyclin/CDK complexes can induce cells in the eyes and wings to proliferate without compromising their ability to undergo terminal differentiation (Buttitta et al., 2007). Profiling of open chromatin during fly wing metamorphosis revealed that only a small subset of genetic loci exhibit changes in chromatin accessibility following direct cell cycle perturbation (Ma et al., 2019). Furthermore, in *C. elegans* development, the overexpression of G1 cyclin/CDK complexes can induce intestinal and body-wall muscle cell proliferation without interfering with the terminally differentiated state of these lineages (Korzelius et al., 2011). These data, along with our results in the invasive AC, suggest that although cell cycle exit is typically linked to the onset of terminal differentiation during normal development, induced proliferation through direct cell cycle perturbation can uncouple this relationship.

We further propose a model in which the loss of negative cell cycle regulators in the AC results in cell cycle entry, but chromatin accessibility is maintained, allowing the pro-invasive GRN to function irrespective of cell cycle state (Fig. 5). Evidence for this model is based on our results, demonstrating that NHR-67 expression is retained in proliferating, invasive ACs. NHR-67 is part of the cell cycle-dependent arm of the GRN and is one of the transcription factors involved in maintaining the post-mitotic state of the invasive AC, as loss of NHR-67 results in proliferating, non-invasive ACs (Matus et al., 2015; Medwig-Kinney et al., 2020). We also examined a downstream effector of the pro-invasive GRN, *zmp-1*, and observed a significant reduction in levels of this MMP gene in proliferating, invasive ACs. However, knockdown of key GRN components results in a complete loss of *zmp-1* transcription (Hwang et al., 2007; Matus et al., 2015; Medwig-Kinney et al., 2020; Rimann and Hajnal, 2007; Sherwood et al., 2005). We suggest that the reduced activity of *zmp-1* in proliferating, invasive ACs may reflect that some aspects of the pro-invasive machinery are affected following cell cycle activation. In the developing fly wing, ectopic activation of the cell cycle results in a reduction in the transcription of genes important for cuticle formation and deposition, which are components of the adult terminal differentiation program (Ma et al., 2019). Notably, cycling wing epithelium cells are still able to secrete cuticle; however, it is thinner than in wild-type non-cycling cells (Ma et al., 2019). Hence, we believe that interference with some aspects of the terminal differentiation program could explain why some, but not all, cycling ACs are able to breach the underlying BM following direct cell cycle perturbation.

With the recent report of the wild-type AC transcriptome (Costa et al., 2023) and advancements in single-cell chromatin and transcription factor profiling (Fabian et al., 2022; Katsanos and Barkoulas, 2022; Katsanos et al., 2021; Yee et al., 2023), the application of these genomic profiling technologies should enable a mechanistic understanding of how the invasive differentiation program can be uncoupled from cell cycle exit. We hope that our results will provide a framework for a better understanding of the interplay between cell cycle exit, terminal differentiation, and cell invasion during normal development and disease states such as cancer metastasis. There is evidence to suggest that at least some tumor cells must alternate between proliferative and invasive states (Kohrman and Matus, 2017; Mondal et al., 2022). However, it is possible that driver mutations in key tumor suppressors such as CKIs or other negative cell cycle regulators can create conditions favorable for tumor cells to maintain their proliferative status while simultaneously adopting an invasive phenotype.

## MATERIALS AND METHODS

### Strains

All strains were grown and maintained following standard procedures (Brenner, 1974). A complete list of strains used in this study is available in Table S1.

### Alleles

The following alleles were used in this study: *qyIs102[fos-1ap::rde-1]* (Hagedorn et al., 2009); **LG I** *bmd156[rps-27p::DHB::2xmKate2]* (Adikes et al., 2020), *bmd294[rps-27p::DHB::2xmTurquoise2]* (this study), and *lin-35(n745)* (Lu and Horvitz, 1998); **LG II** *bmd168[rps-27p::DHB::2xmKate2]* (Adikes et al., 2020), *lin-23(e1883)* (Kipreos et al., 2000), *cki-1(gk132)* (The *C. elegans* Deletion Mutant Consortium, 2012), *cdc-14(he141)* (Saito et al., 2004), *fzr-1(ku298)* (Fay et al., 2002), *cki-2(ok2105)* (The *C. elegans* Deletion Mutant Consortium, 2012), and *rrf-3(pk1426)* (Simmer et al., 2002); **LG IV** *qyIs10[lam-1p::lam-1::GFP]* (Ziel et al., 2009), *qyIs225[cdh-3p::mCherry::moesinABD]* (Matus et al., 2015), and *nhr-67(wy1787[nhr-67::mNeonGreen])* (this study); **LG V** *lag-2(bmd202[lag-2::P2A::H2B::mTurquoise2])* (Medwig-Kinney et al., 2022), *rde-1(ne219)* (Tabara et al., 1999), and *syIs67[zmp-1p::CFP]* (Inoue et al., 2002); **LG X** *lam-2(qy20[lam-2::mNeonGreen])* (Jayadev et al., 2019) and *qyIs24[cdh-3p::mCherry::PLCδPH]* (Ziel et al., 2009).

### Generation of the transgenic bmd294 allele

Both pWZ186 (Addgene plasmid #163641) and a plasmid containing 2xmTurquoise2 were double digested with Bsu36I and NgoMIV to excise 2xmKate2 and 2xmTurquoise2, respectively. pMAM038 (rps-27p::DHB::2xmTurquoise2) was cloned by T4 ligation using the backbone from pWZ186 and 2xmTurquoise2 as an insert. After sequence confirmation, pMAM038 was used as a repair template for insertion into the genome at a safe harbor site on chromosome I corresponding to the MosSCI insertion site, ttTi4348 (Frøkjær-Jensen et al., 2012). pAP082 was used as the sgRNA plasmid for chromosome I insertion by CRISPR/Cas9 (Pani and Goldstein, 2018). Young adults were transformed using standard microinjection techniques, and integrants were identified through the SEC method (Dickinson et al., 2015).

### Generation of the endogenous wy1787 allele

A repair template containing 30xlinker-mNeonGreen, with homology at the 5’ and 3’ ends to the C-terminus of the *nhr-67* locus, was PCR amplified from pJW2171 (Addgene plasmid #163095) and concentrated using a PCR purification kit (Qiagen). 3 µl of 10 µM stock tracRNA (IDT) was incubated with 0.5 µl of a 100 µM crRNA (IDT) for 5 min at 95°C followed by 5 min at 25°C. Immediately following incubation, the tracRNA:cRNA mixture was incubated with 0.5 µl of Alt-R Cas9 protein (IDT) for 10 min at 37°C. The repair template and co-injection marker (pRF6) were combined with the mixture to a final concentration of 200 ng/µl and 50 ng/µl, respectively, for a final injection mix volume of 10 µl. Adult worms possessing no more than a single row of eggs were transformed using standard microinjection techniques. Roller F1 progeny were singled out, and F2 progeny were genotyped for insertions (Paix et al., 2015). The sequence of the PCR primers and guide are available in Tables S2 and S3, respectively.

### RNAi

All RNAi experiments were performed by first synchronizing animals at the L1 larval stage using the bleaching technique (Porta-de-la-Riva et al., 2012). These animals were then fed bacteria expressing the T444T-based RNAi construct for *cki-1* until the mid-L3 larval stage (P6.p four-cell stage), when AC invasion normally occurs (Sherwood and Sternberg, 2003). Control animals were fed bacteria carrying the empty T444T vector.

### Generation of a T444T-based RNAi construct for cki-1

A T444T-based RNAi construct targeting *cki-1* was generated by inserting a 552 bp synthetic DNA fragment, based on the *cki-1* cDNA sequence from WormBase (Davis et al., 2022), into the T444T vector (Addgene plasmid #113081) using the BglII and XhoI restriction sites (Sturm et al., 2018). The synthetic DNA was purchased from IDT as a gBlock with homology arms to T444T for Gibson assembly. The sequence of the synthetic DNA is available in Table S4.

### Image acquisition

Images were acquired using a spinning disk confocal microscope supported by Nobska Imaging. This confocal system consists of a Hamamatsu ORCA EM-CCD camera mounted on an upright Zeiss Axio Imager.A2, equipped with a Borealis-modified Yokogawa CSU-10 spinning disk scanning unit with 6 solid state (405, 440, 488, 514, 561, and 640 nm) lasers and a Zeiss Plan-Apochromat 100x/1.4 oil DIC objective.

For static imaging, animals were anesthetized by placing them into a drop of M9 on a 5% agarose pad containing 7 mM sodium azide and securing them with a coverslip. Time-lapse imaging was performed using a modified version of a previously published protocol (Kelley et al., 2017), as described in Adikes et al. (2020).

### Imaging processing and analysis

The acquired images were processed using ImageJ/Fiji (Schneider et al., 2012). DHB ratios were quantified as previously described (Adikes et al., 2020). NHR-67::mNeonGreen and *zmp-1*::CFP fluorescence were quantified as previously described (Medwig-Kinney et al., 2020). AC invasion was defined as the complete loss of fluorescent BM signal underneath the AC. Plots were generated using Prism software, and figures, including cartoons, were created using a combination of Adobe Photoshop and Illustrator.

### Statistics

A power analysis was performed to determine the number of animals (N) needed per experiment (Pollard et al., 2019) (Table 1). In the text, ‘n’ refers to the number of cells. The figure legends specify the measures of central tendency, error bars, numeric *P*-values, and statistical tests used.

## Acknowledgements

We thank Nicholas Palmisano, Sam Stettnisch, and Abigail Nishimura for their comments and suggestions.

## Data availability

All relevant data can be found within the article and its supplementary information.

## Author contributions

Conceptualization: M.A.Q.M., D.Q.M.; Methodology: M.A.Q.M., D.Q.M.; Validation: M.A.Q.M., C.Z.Z.; Formal analysis: M.A.Q.M.; Investigation: M.A.Q.M., C.Z.Z., F.E.Q.M.; Resources: F.E.Q.M., C.Y., W.Z.; Writing – original draft: M.A.Q.M.; Writing – review and editing: M.A.Q.M., D.Q.M.; Visualization: M.A.Q.M., C.Z.Z., D.Q.M.; Supervision: B.L.M., D.Q.M.; Project administration: M.A.Q.M.; Funding acquisition: K.S., B.L.M., D.Q.M.

---

**Movie 1. Time-lapse imaging confirms that cycling ACs can invade.** A time-lapse of two cycling ACs from pre-invasion to post-invasion is shown. The AC cell membrane, BM, and CDK activity are visualized using an F-actin probe (*cdh-3p*::mCherry::moesinABD), an endogenous *lam-2* translational reporter (LAM-2::mNeonGreen), and a ubiquitous CDK activity sensor (DHB::2xmTurquoise2). The time points were acquired every 5 min for 350 min with a step size of 1 µm. The scale bar represents 5 µm.

