## Supplemental Information for "Cell cycle perturbation uncouples mitotic progression and invasive behavior in a post-mitotic cell"

**Table S1: Strains**

| Name | Genotype | Source |
|---|---|---|
| DQM821 | bmd156[rps-27p::DHB::2xmKate2] I; wild type/mT1 II/III; lag-2(bmd202[lag-2::P2A::H2B::mTurquoise2]) V; lam-2(qy20[lam-2::mNeonGreen]) X | This paper |
| DQM828 | wild type/hT2 I/III; bmd168[rps-27p::DHB::2xmKate2] II; lag-2(bmd202[lag-2::P2A::H2B::mTurquoise2]) V; lam-2(qy20[lam-2::mNeonGreen]) X | This paper |
| DQM995 | bmd156[rps-27p::DHB::2xmKate2] I; cki-2(ok2105) II; lag-2(bmd202[lag-2::P2A::H2B::mTurquoise2]) V; lam-2(qy20[lam-2::mNeonGreen]) X | This paper |
| DQM996 | bmd156[rps-27p::DHB::2xmKate2] I; fzr-1(ku298) unc-4(e120) II; lag-2(bmd202[lag-2::P2A::H2B::mTurquoise2]) V; lam-2(qy20[lam-2::mNeonGreen]) X | This paper |
| DQM1033 | lin-35(n745)/hT2 I; bmd168[rps-27p::DHB::2xmKate2] II; lag-2(bmd202[lag-2::P2A::H2B::mTurquoise2]) V; lam-2(qy20[lam-2::mNeonGreen]) X | This paper |
| DQM1098 | bmd156[rps-27p::DHB::2xmKate2] I; wild type/mIn1 II; lag-2(bmd202[lag-2::P2A::H2B::mTurquoise2]) V; lam-2(qy20[lam-2::mNeonGreen]) X | This paper |
| DQM1110 | bmd156[rps-27p::DHB::2xmKate2] I; cdc-14(he141) II; lag-2(bmd202[lag-2::P2A::H2B::mTurquoise2]) V; lam-2(qy20[lam-2::mNeonGreen]) X | This paper |
| DQM1111 | bmd156[rps-27p::DHB::2xmKate2] I; lin-23(e1883)/mIn1 II; lag-2(bmd202[lag-2::P2A::H2B::mTurquoise2]) V; lam-2(qy20[lam-2::mNeonGreen]) X | This paper |
| DQM1174 | bmd294[rps-27p::DHB::2xmTurquoise2] I; fzr-1(ku298) unc-4(e120) II; qyIs225[cdh-3p::mCherry::moesinABD] V; lam-2(qy20[lam-2::mNeonGreen]) X | This paper |
| DQM1207 | bmd156[rps-27p::DHB::2xmKate2] I; rrf-3(pk1426) II; lag-2(bmd202[lag-2::P2A::H2B::mTurquoise2]) V; lam-2(qy20[lam-2::mNeonGreen]) X | This paper |
| DQM1316 | bmd156[rps-27p::DHB::2xmKate2] I; fzr-1(ku298) unc-4(e120) II; nhr-67(wy1787[nhr-67::mNeonGreen]) IV; lag-2(bmd202[lag-2::P2A::H2B::mTurquoise2]) V; lam-2(qy20[lam-2::mNeonGreen]) X | This paper |
| DQM1317 | bmd156[rps-27p::DHB::2xmKate2] I; fzr-1(ku298) unc-4(e120) II; syIs67[zmp-1::pes-10::CFP] V; lam-2(qy20[lam-2::mNeonGreen]) X | This paper |
| NK1316 | qyIs102[fos-1ap::rde-1]; rrf-3(pk1426) II; qyIs10[lam-1p::lam-1::GFP] IV; rde-1(ne219) V; qyIs24[cdh-3p::mCherry::PLCδPH] X | Morrissey et al., 2014 |
| VC170 | cki-1(gk132)/mIn1 II | CGC |

**Table S2: Primers**

| Name | Sequence (5' – 3') | Type | Amplicon | Template |
|---|---|---|---|---|
| CY584 | TTCCTCAACTCAGCCTTCA<br>TCGGCATCATCCCCTTCC<br>TCTTCAAGACCACGTCATT<br>CGATTCGATCAATAACTG<br>AATTATTATCAATTCAAGA<br>AGAgGAaAGcGTgAAcGTg<br>GAgGAaGTGggtggcggtggat<br>cgggaggaggaggttc | Forward | nhr-67::mNeonGreen | pJW2171 |
| CY729 | ttttgaaagtataaatcaaaatccaatg<br>aaaatccgggcgaggcaagaaacg<br>gagtgaaaaagacaatgaaaagag<br>atacttagaataattgagttactgtatat<br>gaaacatgaatttactatTTACTTG<br>TAGAGCTCGTCCATTCCC<br>AT | Reverse | nhr-67::mNeonGreen | pJW2171 |

**Table S3: Guides**

| Locus | Sequence (5' – 3') | Description |
|---|---|---|
| *nhr-67* | AGAGAGTGTTAATGTTGAAGAGG | Located upstream of the stop codon |

**Table S4: gBlocks**

| Gene | Sequence (5' – 3') |
|---|---|
| *cki-1* | tcactatagggagaccggcaATGTCTTCTGCTCGTCGTTGCCTTTTCGGTCGT<br>CCGACGCCCGAGCAACGCTCCAGGACTCGAATTTGGCTTGAAGATG<br>CTGTTAAGCGCATGCGCCAGGAAGAAAGCCAGAAATGGGGATTCGA<br>CTTTGAACTGGAGACTCCCCTCCCAAGCTCTGCTGGATTCGTTTATG<br>AAGTTATTCCAGAGAATTGTGTTCCGGAGTTCTACAGAACCAAAGTT<br>CTCACTGTCAGAACCACATGCTCATCGCTGGACATCAGCTCAACGA<br>CTTTGACTCCATTGAGCTCTCCGAGCACATCTGATAAGGAGGAGCC<br>CTCGCTGATGGATCCCAACAGCTCGTTCGAAGATGAAGAGGAACCG<br>AAGAAGTGGCAATTCAGAGAGCCACCAACTCCACGGAAGACCCCAA<br>CAAAGCGTCAGCAGAAGATGACCGACTTCATGGCAGTTTCCCGTAA<br>GAAGAATTCGTTGTCTCCAAACAAGCTGTCTCCGGTGAATGTGATCT<br>TCACTCCAAAATCTCGTCGTCCAACGATCAGAACTCGATCTTCATGC<br>TCTCCATACgggggggcccggtacccaat |

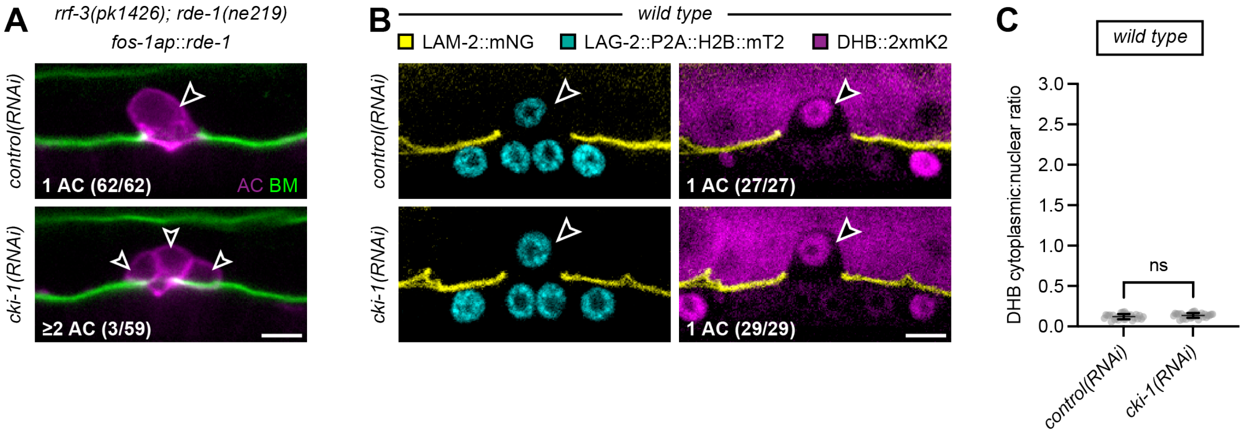


**Figure S1. AC proliferation is rarely observed upon uterine-specific depletion of CKI-1.** (A) Confocal images merging *cdh-3p*::mCherry::*PLCδPH* and *lam-1p*::LAM-1::GFP from the P6.p 4-cell stage are shown. Arrowheads indicate the position of the AC(s) in a uterine-specific RNAi strain (*rrf-3(pk1426); rde-1(ne219); fos-1ap::rde-1*) following treatment with *control(RNAi)* and *cki-1(RNAi)*. The scale bar represents 5 µm. (B) Confocal images merging LAM-2::mNeonGreen with LAG-2::P2A::H2B::mTurquoise2 (left) and DHB::2xmKate2 (right) from the P6.p 4-cell stage are shown. Arrowheads indicate the position of the AC in wild-type animals following *control(RNAi)* and *cki-1(RNAi)* treatment. The scale bar represents 5 µm. (C) Scatter plot displays the mean and SD of DHB::2xmKate2 ratios in the AC of wild-type animals following *control(RNAi)* and *cki-1(RNAi)* treatment (N ≥ 27 animals per treatment). Statistical significance was determined by an unpaired t test (ns).

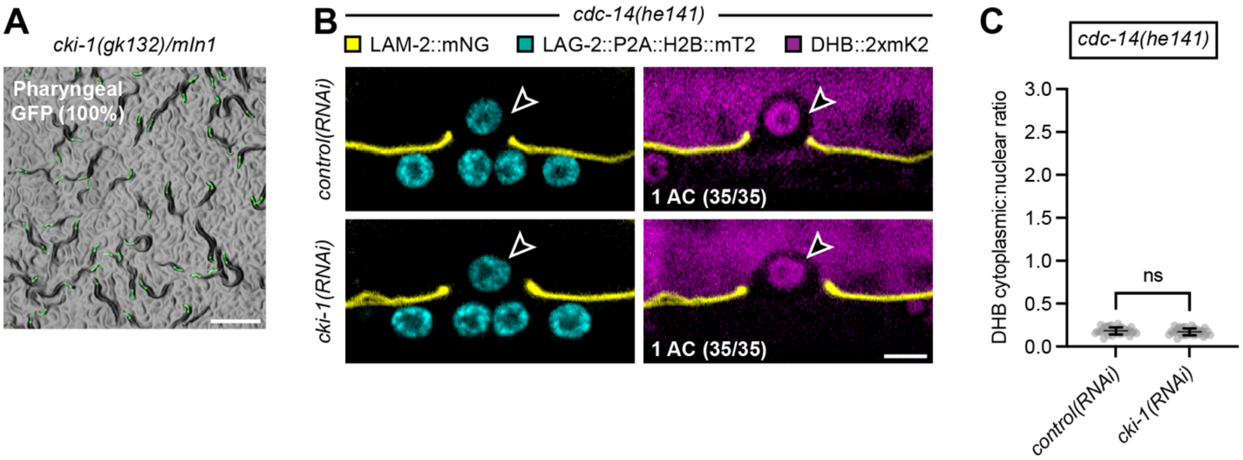


**Figure S2. CDC-14 is not an essential regulator of CKI-1 in the AC.** (A) A stereoscopic image merging brightfield and *myo-2p*::GFP is shown. At the mid-L3 larval stage, 100% of progeny from *cki-1(gk132)/mIn1* mothers exhibit pharyngeal GFP expression, as *cki-1(gk132)* homozygotes are embryonic lethal. The scale bar represents 500 µm. (B) Confocal images merging LAM-2::mNeonGreen with LAG-2::P2A::H2B::mTurquoise2 (left) and DHB::2xmKate2 (right) from the P6.p 4-cell stage are shown. Arrowheads indicate the position of the AC in *cdc-14(he141)* animals following *control(RNAi)* and *cki-1(RNAi)* treatment. The scale bar represents 5 µm. (C) Scatter plot displays the mean and SD of DHB::2xmKate2 ratios in the AC of *cdc-14(he141)* animals following *control(RNAi)* and *cki-1(RNAi)* treatment (N = 35 animals per treatment). Statistical significance was determined by an unpaired t test (ns).

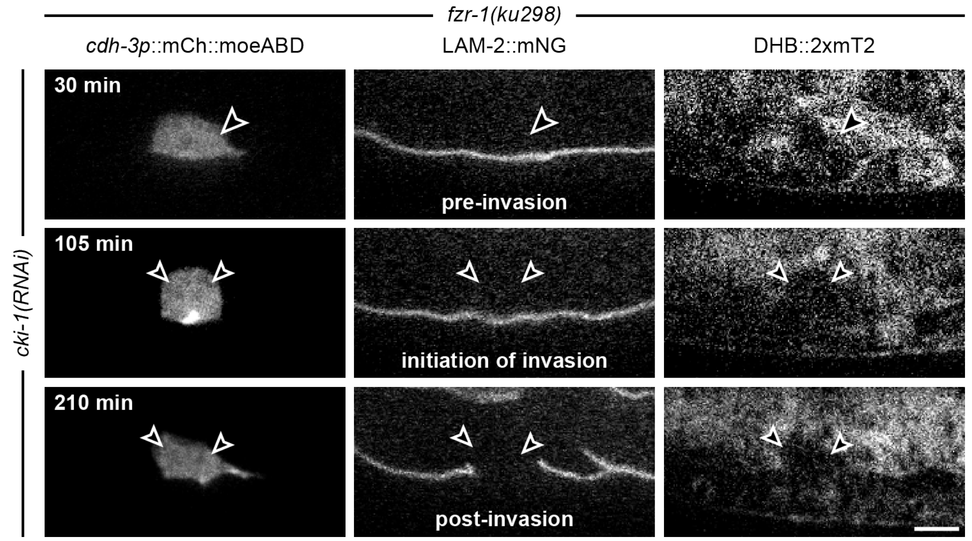


**Figure S3. Additional evidence confirming that the maintenance of the post-mitotic state is not required for AC invasion.** Time-lapse images of *cdh-3p*::mCherry::moesinABD (left), LAM-2::mNeonGreen (middle), and DHB::2xmTurquoise2 (right) from pre-AC invasion to post-AC invasion are shown. Arrowheads indicate the position of a cycling AC (top) that subsequently divides, re-enters the cell cycle, and invades (middle and bottom) in a *fzr-1(ku298); cki-1(RNAi)* animal. At the initiation of invasion time-point, the DHB::2xmTurquoise2 ratio is 1.23 for the AC on the left and 1.31 for the AC on the right. The scale bar represents 5 μm.
